# Validated Preclinical Murine Model for Therapeutic Testing against Multidrug Resistant *Pseudomonas aeruginosa*

**DOI:** 10.1101/2022.07.05.498916

**Authors:** Jonathan M. Warawa, Xiaoxian Duan, Charles D. Anderson, Julie B. Sotsky, Daniel E. Cramer, Tia L. Pfeffer, Haixun Guo, Robert S. Adcock, Alexander J. Lepak, David R. Andes, Stacey A. Slone, Arnold J. Stromberg, Jon D. Gabbard, William E. Severson, Matthew B. Lawrenz

## Abstract

The rise in infections caused by antibiotic resistant bacteria is outpacing the development of new antibiotics. The ESKAPE pathogens (*Enterococcus faecium, Staphylococcus aureus, Klebsiella pneumoniae, Acinetobacter baumannii, Pseudomonas aeruginosa*, and *Enterobacter* species) are a group of clinically important bacteria that have developed resistance to multiple antibiotics and are commonly referred to as multidrug resistant (MDR). The medical and research communities have recognized that without new antimicrobials, infections by MDR bacteria will soon become a leading cause of morbidity and mortality. Therefore, there is an ever growing need to expedite the development of novel antimicrobials to combat these infections. Toward this end, we set out to refine an existing murine model of pulmonary *Pseudomonas aeruginosa* infection to generate a robust preclinical tool that can be used to rapidly and accurately predict novel antimicrobial efficacy. This refinement was achieved by characterizing the virulence of a panel of genetically diverse MDR *P. aeruginosa* strains in this model, both by LD_50_ analysis and natural history studies. Further, we defined two antibiotic regimens (aztreonam and amikacin) that can be used a comparators during the future evaluation of novel antimicrobials, and validated that the model can effectively differentiate between successful and unsuccessful treatment as predicted by in vitro inhibitory data. This validated model represents an important tool in our arsenal to develop new therapies to combat MDR *P. aeruginosa*, with the ability to provide rapid preclinical evaluation of novel antimicrobials that can also serve to support data from clinical studies during the investigational drug development process.

## Introduction

It is estimated that ≥1,000,000 people worldwide die each year due to infections by antibiotic-resistant bacteria [1]. Based on the current rate of antibiotic resistance observed in the clinic, and without a concerted effort to identify and develop new antibiotics, the United Nations Interagency Coordinating Group on Antimicrobial Resistance predicted mortality due to antibiotic-resistant bacteria could increase by several million by 2050 [2]. The ESKAPE pathogens (*Enterococcus faecium, Staphylococcus aureus, Klebsiella pneumoniae, Acinetobacter baumannii, Pseudomonas aeruginosa*, and *Enterobacter* species) have been recognized by the CDC as the leading cause of nosocomial infections throughout the world and have all acquired resistance to multiple antibiotics. *P. aeruginosa* is an opportunistic pathogen that can cause infection of the skin, eyes, urinary tract, and lungs, and is especially pathogenic for immunocompromised individuals. On its own, *P. aeruginosa* is estimated to be responsible for up to 20% of nosocomial infections within intensive care units in the USA and European Union, of which >35% of isolates are resistant to multiple antibiotics[3-5]. Infections with multidrug resistant (MDR) *P. aeruginosa* are associated with significant increases in mortality, particularly in patients with respiratory-associated bacteremia [6].

To combat these infections, we need to identify and develop new antimicrobials against the ESKAPE pathogens. Toward this goal is the development of robust preclinical models that can quickly and accurately predict the efficacy of novel antimicrobials in humans. The mouse is the most widely used model for preclinical testing of antimicrobial investigational drugs (INDs). The main driving factors for this are 1) a large amount of data available on the pathogenesis of microbes in the mouse, 2) inbred mouse strains limit variation/noise due to genetic drift, and 3) the low cost to purchase, breed, and house mice allows for increased group sizes and statistical power to overcome experimental noise often associated with preclinical models. Experimental noise is considered a major contributor to past failures in translation of preclinical results into the clinic [7]. Therefore, this third driving factor is critically important for successful preclinical testing. Importantly, the application of pharmacokinetics/ pharmacodynamics analysis of antimicrobials in mice allow for the identification of driver indices that are independent of host metabolism and can be used to bridge preclinical data to human application [8]. Therefore, preclinical data from mice on the efficacy of antimicrobials is significantly more predictive of success in the clinic than other classes of drugs. However, for the mouse to be an effective preclinical model, it should be well characterized to recapitulate human disease and validated to respond to current antimicrobials as expected with a variety of strains representing the potential diversity of isolates seen in the clinic.

We have previously developed a lethal respiratory model of MDR *P. aeruginosa* infection in leukopenic mice for the purpose of testing novel therapeutics [9]. This model recapitulates human disease, with bacterial proliferation in the lungs, acute inflammation, and development of pneumonia [9-11]. Using this model, we have successfully evaluated efficacy for numerous therapies either in monotherapy or as a combination therapy with a representative carbapenem antibiotic, meropenem [9, 12, 13]. However, this model has limitations, including characterization against only one strain of *P. aeruginosa* and a lack of robust comparative antibiotics to gauge the success of novel INDs. Therefore, to improve our current model for preclinical screening of antimicrobials against *P. aeruginosa*, we validated our infection model with four additional strains of MDR *P. aeruginosa* better representing the genetic diversity of clinical isolates. We also fully characterized two antibiotics under success and failure scenarios so that they can be used as comparator drugs to gauge the efficacy of novel antimicrobials against acute pulmonary infection by *P. aeruginosa*.

## Results

### Characterization of P. aeruginosa strains in the leukopenic mouse model – Identification of the lethal dose 50

Our first step in enhancing the predictive ability of the mouse model we previous developed for preclinical screening of antimicrobials against *P. aeruginosa* infection [9] was to characterize the virulence of different strains of MDR *P. aeruginosa* within this model. Toward this goal, we chose to characterize four *P. aeruginosa* strains that are widely available to the research community through the FDA-CDC Antimicrobial Resistance Isolate Bank *P. aeruginosa* Panel (https://www.cdc.gov/drugresistance/resistance-bank/currentlyavailable.html). From this panel, the strains were chosen because their genomes have been sequenced, they have different antibiotic resistance profiles, including different known mechanisms to β-lactams, and they display different degrees of resistance to amikacin and aztreonam, which can be used as comparators in future studies (Table 1).

**Table 1.** *P. aeruginosa* strains characterized in mouse model.

| Table 1. *P. aeruginosa* strains characterized in mouse model. |  |  |  |  |  |
| --- | --- | --- | --- | --- | --- |
| Strain | Antibiotic Phenotype* |  |  | Known Resistance | Genome Accession # |
|  | Resistant | Sensitive | Inter. |  |  |
| 0230 | **Ami**, Cefe, Ceft, Cip, Dor, Gen, Imi, Lev, Mer, Pip, Tob | **Azt**, Col, Pol | -- | VIM | SAMN04901620 |
| 0231 | **Azt**, Cefe, Ceft, Cip, Dor, Gen, Imi, Lev, Mer, Pip, Tob | **Ami**, Col, Pol | -- | KPC | SAMN04901621 |
| 0241 | Cefe, Ceft, Cip, Dor, Gen, Imi, Lev, Mer, Pip, Tob | Col, Pol | **Ami**,<br>**Azt** | IMP | SAMN04901631 |
| 0246 | **Ami**, **Azt**, Cefe, Ceft, Cip, Dor, Gen, Imi, Lev, Mer, Pip, Tob | Col, Pol | -- | NDM | SAMN04901636 |
| *Amikacin (Ami); Aztreonam (Azt); Cefepime (Cefe); Ceftazidime (Ceft); Ciprofloxacin (Cip); Colistin (Col); Doripenem (Dor); Gentamicin (Gen); Imipenem (Imi); Levofloxacin (Lev); Meropenem (Mer); Piperacillin/Tazobactam (Pip); Polymyxin B (Pol); Tobramycin (Tob) |  |  |  |  |  |

Because no virulence data is available for the isolates within the FDA-CDC Antimicrobial Resistance Isolate Bank *P. aeruginosa* Panel, we first sought to identify the number of bacteria required to cause lethal infection in 50% of infected animals (i.e., the Lethal Dose 50 or LD_50_) for each strain in cyclophosphamide-induced leukopenia mice [9]. Male and female mice were infected by direct lung instillation with escalating doses of the four MDR *P. aeruginosa* strains [9, 14]. Mice were monitored for the development of moribund disease at 8 h intervals. Mice that met predefined endpoint criteria were humanely euthanized and scored as succumbing to disease 8 h later. All four strains were able to establish lethal infection in leukopenic mice, but we observed differences in the LD_50_ between the strains. Strains 0230 and 0231 were the most virulent in the mouse, with >50% mortality achieved with only 10^2^ CFU challenge (Fig. 1A-B). The 0246 strain was slightly less virulent, as >50% of animals succumbed to infection at the 10^3^ CFU dose (Fig. 1C). In contrast, strain 0241 was significantly less virulent than the other three strains, requiring 10^6^ CFU to achieve >50% mortality (Fig. 1C). For all strains tested, sex did not have a significant impact on the resistance of the host to lethal infection. Finally, at inoculums resulting in ∼50% mortality, we observed similar mean times to death for all strains, ranging from 44-56 h. From these survival curves we calculated the LD_50_ for strains 0230, 0231, 0246 and 0241 as 1.30, 1.59, 2.78, and 5.61 log CFU, respectively.

**Fig. 1.**
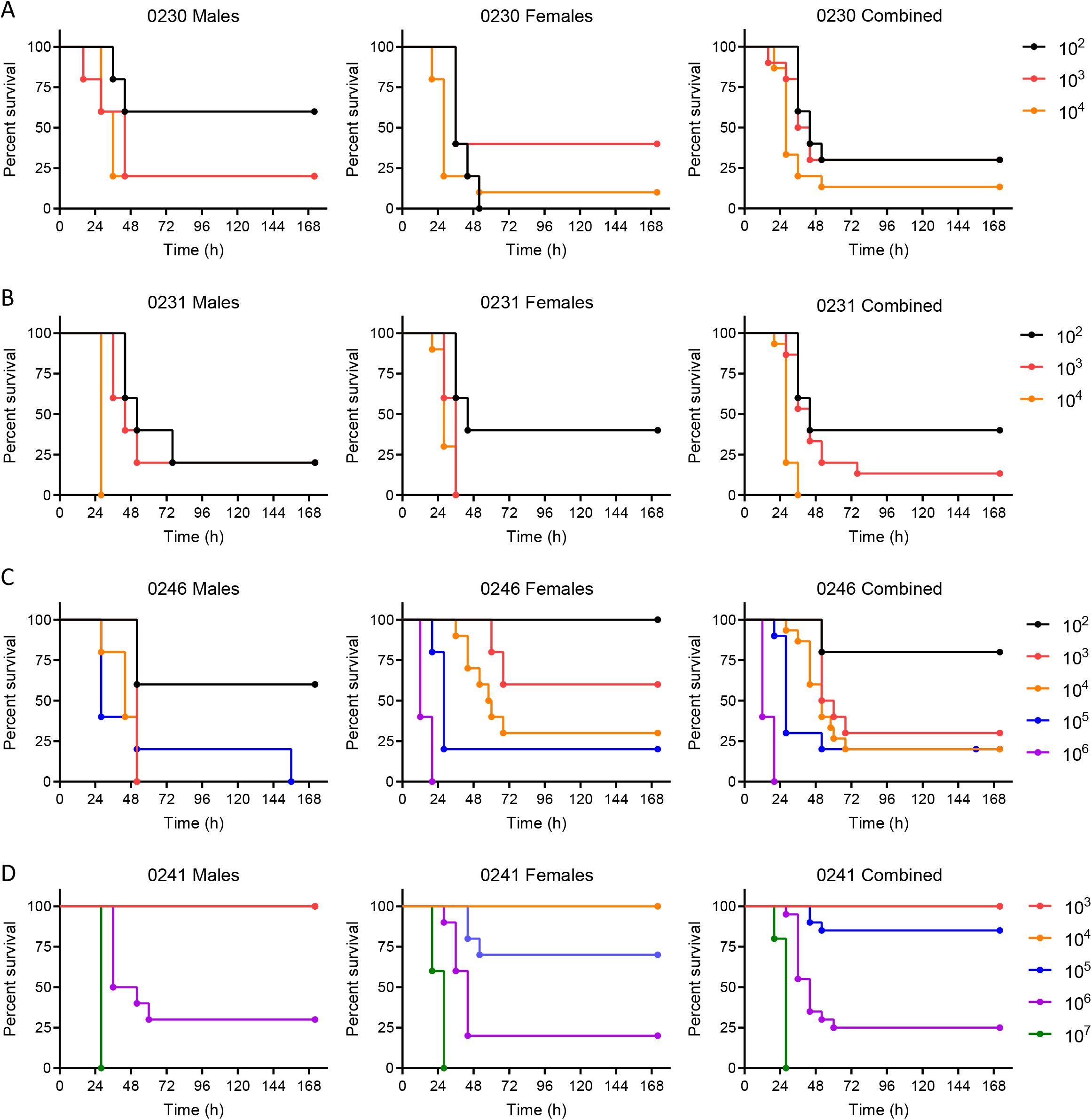
Survival of mice infected with different strains of *P. aeruginosa*. Equal numbers of male and female mice (n=10) were infected with indicated CFU of *P. aeruginosa* strains (A) 0230, (B) 0231, (C) 0246, or (D) 0241 and monitored every 8 h for the development of moribund disease. Animals that met predetermined endpoint criteria were scored as succumbing to infection 8 h later.

### Characterization of P. aeruginosa strains in the leukopenic mouse model – Natural history of infection

Once we established that each strain was able to cause lethal infection, we next sought to characterize the natural history of the infection in the leukopenic mouse model. Male and female mice were infected by direct instillation into the lungs with 10x the LD_50_ of each of the *P. aeruginosa* strains. At 3, 6, 12, and 21 h host temperatures were measured, and subsets of mice were humanely euthanized to harvest tissues for bacterial enumeration and histological analysis. 21 h was chosen as the last timepoint to minimize the number of animals reaching endpoint criteria prior to a designated time point and resulting in a need to exclude said animals from subsequent analysis. In general, mice that received the more virulent strains 0230, 0231, and 0246, and were inoculated with 3-4 log CFU of bacteria, followed very similar courses of disease. At 3 h post-infection, ∼3.0 log CFU of bacteria were recovered from the lungs of infected mice. Similar numbers were recovered at 6 h post-infection, but by 12 h post-infection, bacterial numbers increased by ∼1.5 log. By 21 h post-infection, mean bacterial numbers were >6 log CFU for all three strains (Fig. 2). Bacterial dissemination from the lungs to the blood as indicated by spleen colonization was not detected prior to 21 h for any of the three strains (Fig. 2). Significant changes in the host response to infection, measured by hypothermia and lung inflammation, was also not observed in 0230, 0231, and 0246 infected mice until the 21 h post-infection (Fig. 3).

**Fig. 2.**
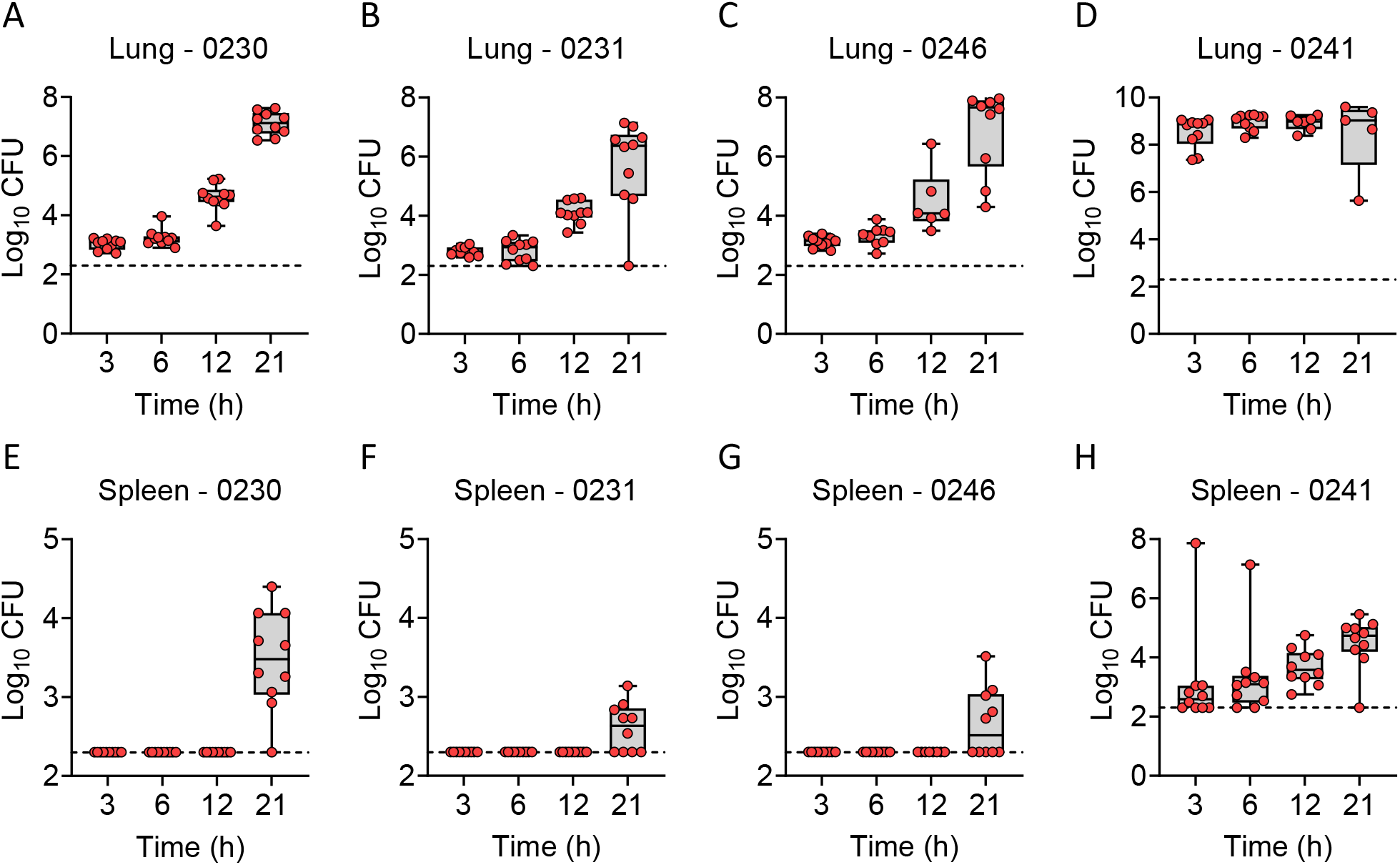
Bacterial enumeration from tissues of infected mice. Equal numbers of male and female mice (n=10) were infected with 10x the LD_50_ of indicated *P. aeruginosa* strains. At 3, 6, 12, and 21 h, mice were euthanized and bacterial numbers were enumerated from the lungs (A-D) or spleens (E-H). Each circle represents data from an individual mouse. The dotted line indicates the limit of detection.

**Fig. 3.**
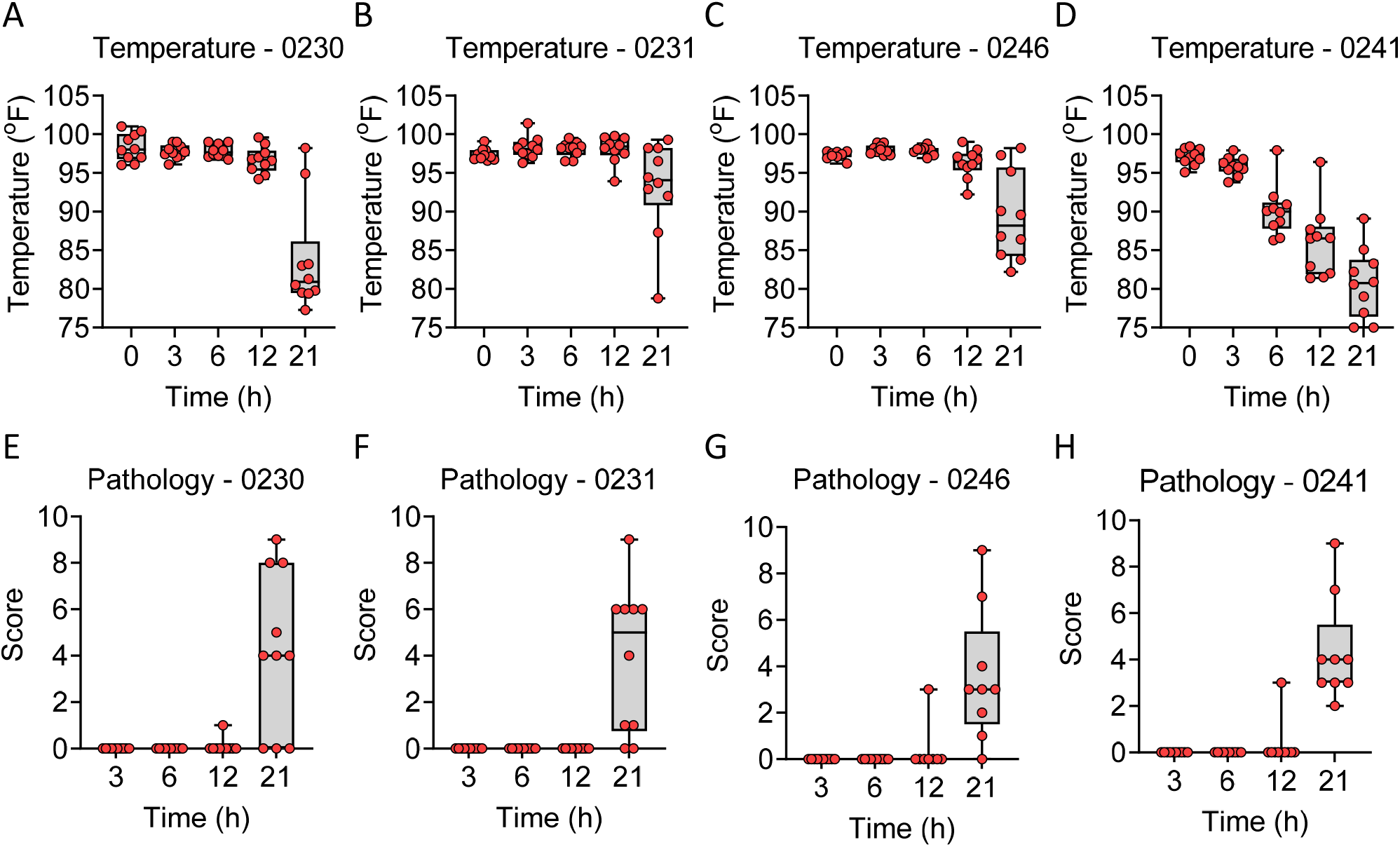
Host response during *P. aeruginosa* infection. Equal numbers of male and female mice (n=10) were infected with 10x the LD_50_ of indicated *P. aeruginosa* strains. (A-D) Temperature measurements at indicated times. (E-H) At 3, 6, 12, and 21 h, mice were euthanized, and a cross section of the lungs was excised, stained with H&E, and inflammation and pathology was assessed blindly. Each circle represents data from an individual mouse.

Strain 0241 being less virulent than the other three strains required a significant higher inoculum to establish a lethal disease (10x LD_50_ = 6.6 log CFU). As such, significantly higher bacterial numbers were recovered from the lungs by 3 h post-infection (Fig. 2D). Bacterial burdens did not change over the 21 h period of observation. However, the elevated bacterial numbers in the lungs resulted in bacterial dissemination to the bloodstream and spleen within 3 h of inoculation (Fig. 2H). This earlier dissemination directly correlated with earlier development of hypothermia in the animals infected with strain 0241 (Fig. 3D). However, despite increased bacterial burdens at earlier time points, lung inflammation and pathology were still not observed 21 h post-infection (Fig. 3H). Together these data establish that the leukopenic mouse model is amenable to infection by a variety of different MDR *P. aeruginosa* strains and established parameters of infection that can be used to measure subsequent drug efficacy, including bacterial replication, dissemination, onset of hypothermia, and lung pathology.

### Characterization of comparator antibiotics - Pharmacokinetics of aztreonam and amikacin

Our second goal in improving the predictive ability of the mouse model for preclinical screening of antimicrobials against *P. aeruginosa* infection was to establish an antibiotic that could be used as a comparator to gauge the efficacy of novel INDs. Part of the criteria during the selection of the panel of *P. aeruginosa* clinical isolates characterized above was differential susceptibility to an antibiotic that would allow for us to demonstrate both success and failure scenarios in the model. As such, strains 0230, 0231, 0246, and 0241 have different susceptibility to two antibiotics, aztreonam and amikacin (Table 1). Based on these phenotypes, we next sought to establish pharmacokinetic parameters for both antibiotics in the leukopenic mouse model during *P. aeruginosa* infection. Male and female mice were infected by direct instillation into the lung with 10x the LD_50_ of either strain 0230 (for mice receiving aztreonam) or strain 0231 (for mice receiving amikacin). At 3 h post-infection, mice received escalating doses of antibiotics and plasma was collected at 5, 10, 15, 30, 60 and 120 min post-antibiotic administration. Antibiotic concentrations in plasma were quantified and analytically assessed by high performance liquid chromatography (HPLC) (Fig. 4, Tables 2 and 3). Pharmacokinetic constants were calculated using a time ordered dataset generated from the pooled pharmacokinetic data.

**Fig. 4.**
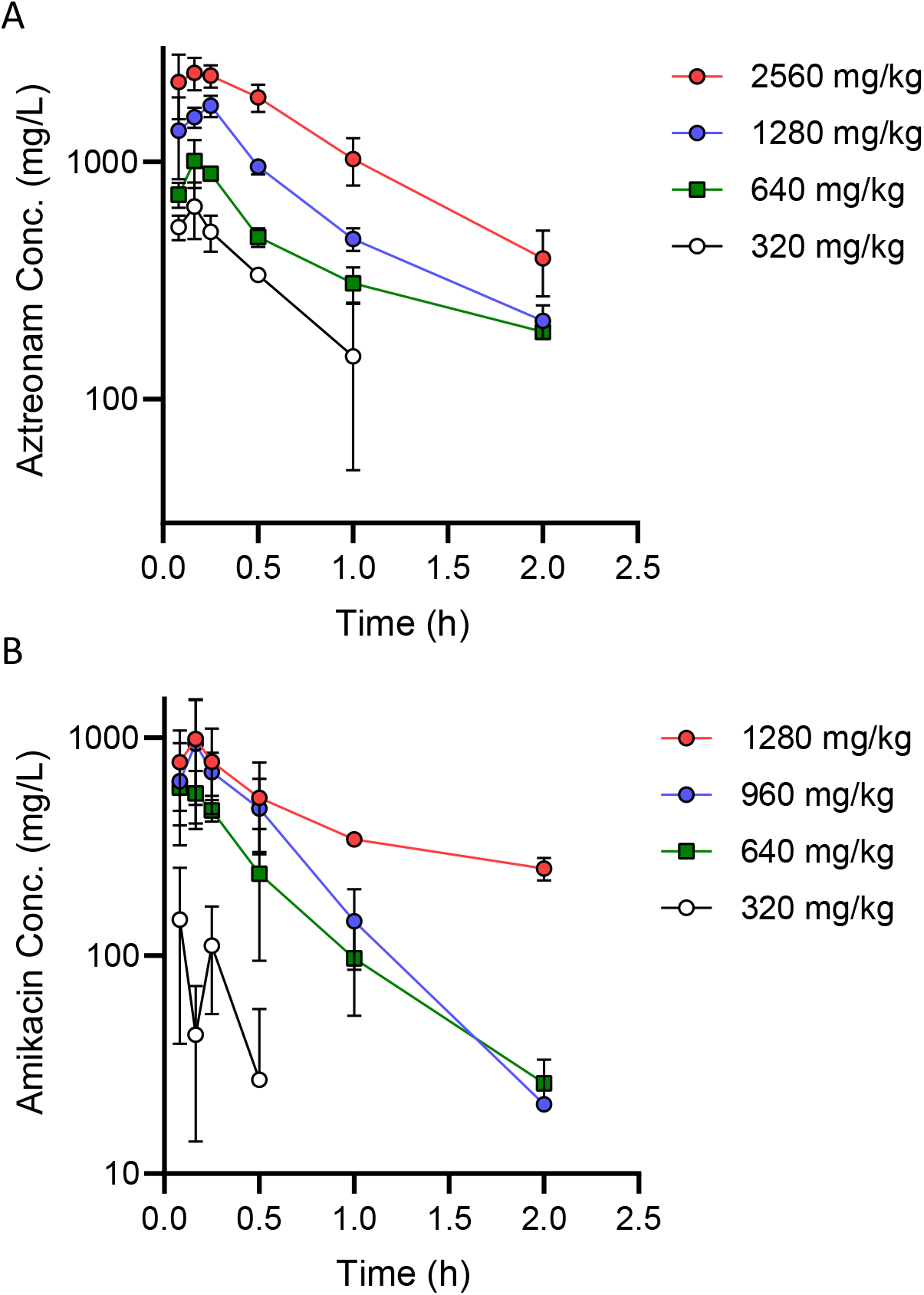
Concentration of antibiotics in plasma. Mice (n=4) were administered a single dose of (A) aztreonam or (B) amikacin and at indicted time points drug concentration was determined for plasma.

**Table 2.** Aztreonam plasma concentrations over time.

| Time point (min) | 320 mg/kg |  | 640 mg/kg |  | 1280 mg/kg |  | 2560 mg/kg |  |
| --- | --- | --- | --- | --- | --- | --- | --- | --- |
|  | Mean (ug/mL) | St. dev | Mean (ug/mL) | St. dev | Mean (ug/mL) | St. dev | Mean (ug/mL) | St. dev |
| 5 | 530.3 | 63 | 727.6 | 87.4 | 1353.8 | 511 | 2171 | 659 |
| 10 | 646.4 | 173 | 1005.2 | 229.5 | 1542 | 151 | 2372 | 374 |
| 15 | 506.2 | 87.5 | 889.3 | 45.7 | 1720 | 180 | 2303 | 249 |
| 30 | 333.3 | 9.8 | 481 | 43.1 | 953.1 | 70 | 1867 | 248 |
| 60 | 151.2 | 101.1 | 307.1 | 51.8 | 472.3 | 51.3 | 1026 | 233 |
| 120 | BLQ | -- | 191.9 | 7.5 | 213.7 | 34.5 | 392 | 121 |

**Table 3.** Amikacin plasma concentrations over time.

| Time point (min) | 320 mg/kg |  | 640 mg/kg |  | 960 mg/kg |  | 1280 mg/kg |  |
| --- | --- | --- | --- | --- | --- | --- | --- | --- |
|  | Mean (ug/mL) | St. dev | Mean (ug/mL) | St. dev | Mean (ug/mL) | St. dev | Mean (ug/mL) | St. dev |
| 5 | 145.97 | 106.47 | 589.70 | 195.16 | 630.96 | 310.37 | 769.89 | 307.99 |
| 10 | 43.31 | 29.23 | 553.54 | 149.92 | 938.60 | 558.66 | 987.57 | 496.23 |
| 15 | 110.85 | 56.91 | 463.84 | 52.67 | 695.31 | 155.22 | 772.94 | 326.85 |
| 30 | 26.94 | 29.94 | 237.38 | 142.52 | 472.03 | 173.56 | 529.99 | 238.09 |
| 60 | BLQ | -- | 97.20 | 44.14 | 143.93 | 57.97 | 340.95 | 23.33 |
| 120 | BLQ | -- | 25.92 | -- | 20.80 | 12.57 | 250.90 | 29.55 |

Using these data, we calculated dosing regimens for mice that would closely mimic those parameters achieved in the clinic for humans. For aztreonam, time the drug is above the minimal inhibitory concentration (T>MIC) is the primary driver of success. Using human clinical data based on administration of 2 gm of aztreonam intravenously Q8h as our target [15], we calculated the T>MIC in mice receiving various regimens of aztreonam via subcutaneous injection (Table 4). Based on our pharmacokinetics data, 640 mg/kg/6h SC in the mouse is predicted to achieve a free Cmax slightly higher than humans but will achieve T>MIC exposures against organisms (susceptible and resistant) that would be predicted to be therapeutically successful for a susceptible organism (MIC=4 μg/ml), marginal for a resistant strain (MIC=32 μg/ml), and unsuccessful for a highly resistant strain (MIC>64 μg/ml).

**Table 4.** Time above the MIC for aztreonam in mice.

| **Human** |  |  |  |  |
| --- | --- | --- | --- | --- |
| **Dose**<br>**(mg/kg)** | **Dosing**<br>**Interval (h)** | **Free Drug % Time**<br>**above MIC (4 mg/L)** | **Free Drug % Time**<br>**above MIC (32 mg/L)** | **Free Drug % Time**<br>**above MIC (>64 mg/L)** |
| 2 g | 8 | 97 | 38 | 0 |
| **Mouse** |  |  |  |  |
| **Dose**<br>**(mg/kg)** | **Dosing**<br>**Interval (h)** | **Free Drug % Time**<br>**above MIC (4 mg/L)** | **Free Drug % Time**<br>**above MIC (32 mg/L)** | **Free Drug % Time**<br>**above MIC (>64 mg/L)** |
| 320 | 4 | 51 | 21 | 0 |
| 640 | 4 | 100 | 45 | 5 |
| 1280 | 4 | 96 | 50 | 19 |
| 2560 | 4 | 100 | 66 | 32 |
| 320 | 6 | 34 | 14 | 0 |
| 640 | 6 | 69 | 30 | 3 |
| 1280 | 6 | 64 | 33 | 12 |
| 2560 | 6 | 78 | 44 | 22 |
| 320 | 8 | 26 | 10 | 0 |
| 640 | 8 | 52 | 22 | 2 |
| 1280 | 8 | 48 | 25 | 9 |
| 2560 | 8 | 59 | 33 | 16 |
| 320 | 12 | 17 | 7 | 0 |
| 640 | 12 | 32 | 15 | 1 |
| 1280 | 12 | 35 | 17 | 6 |
| 2560 | 12 | 39 | 22 | 11 |

**Table 5.** Calculated AUC0-24 and Cmax for amikacin.

| **Human Amikacin dose** |  | **Human AUC<sub>0-24</sub>** | **Human Cmax** |
| --- | --- | --- | --- |
| Q24 | 15 mg/kg | 370 mg*h/L | 65 mg/L |
| **Mouse Amikacin dose** |  | **Mouse AUC<sub>0-24</sub>** | **Mouse Cmax** |
| Q24 | 605 mg/kg | 370 mg*h/L | 541 mg/L |
| Q24 | 142 mg/kg | 20 mg*h/L | 65 mg/L |
| Q12 | 385 mg/kg | 370 mg*h/L | 236 mg/L |
| Q12 | 142 mg/kg | 40 mg*h/L | 65 mg/L |
| Q6 | 293 mg/kg | 370 mg*h/L | 109 mg/L |
| Q6 | 142 mg/kg | 80 mg*h/L | 65 mg/L |

For amikacin, the area under the curve (AUC_0-24_) is the primary driver of success. Using human clinical data based on the 15 mg/kg of amikacin administered to patients Q24h as our target [16, 17], we calculated dosing regimens of amikacin via subcutaneous injection in mice that matched the AUC_0-24_ and Cmax seen in human patients (Table 4). Treating mice Q24h with a dose of 605 mg/kg would match the AUC_0-24_ but exceed the Cmax by 8.3-fold. Similarly, treating mice Q12h with a dose of 385 mg/kg would also match the AUC_0-24_ but exceed the Cmax by 3.6-fold. Finally, treating mice Q6h with a dose of 293 mg/kg would match the AUC_0-24_ but only exceed the Cmax by 1.7-fold. Based on these data, we predicted a drugging regimen of 640 mg/kg/6h for aztreonam and 293 mg/kg/6h for amikacin will best represent a humanized dose of each drug in the leukopenic mouse model.

### Validation of humanized doses of antibiotics - Bacterial reduction

Having identified humanized dosing regimens for aztreonam and amikacin, we next sought to validate these regimens by demonstrating both success and failure scenarios in the leukopenic mouse model. Toward this goal, equal numbers of male and female mice were infected by direct instillation into the lungs with 10x the LD_50_ of *P. aeruginosa* strain 0230 (Azt^S^, Ami^R^) or 0231 (Azt^R^, Ami^S^). At 6 h post-infection, mice were administered 640 mg/kg aztreonam, 293 mg/kg amikacin, or vehicle only (PBS). Drug administration was repeated every 6 h. At each administration, mouse temperature was also measured. At 21 h mice were euthanized and tissues were harvested for bacterial enumeration and histological analysis. Results were compared to tissues harvested from a subset of mice 6 h post-infection, representing the baseline values prior to the first administration of antibiotic.

As predicted by the in vitro MIC determinations, mice infected with *P. aeruginosa* 0230 (Azt^S^, Ami^R^) began developed sings of infection, including hypothermia, at 12 h post-infection when treated with PBS or amikacin, but not if the animals received aztreonam (Fig. 5A). While low numbers of bacteria were recovered from the lungs in some of the aztreonam-treated animals, the bacterial burden was significantly higher in the lungs of the PBS- and amikacin-treated mice, representing a difference of >3.0 log CFU between the groups (Fig. 5B; p>0.0001). Moreover, no bacteria were recovered from the spleens of aztreonam-treated mice, while mice treated with PBS or amikacin had disseminated infection represented by significantly higher bacterial burden in the spleens (Fig. 5C; p>0.01). Finally, while mild lung pathology was observed in aztreonam-treated mice, pathology scores were significantly higher in animals treated with PBS or amikacin. Similar trends were observed for mice infected with *P. aeruginosa* 0231 (Azt^R^, Ami^S^) except health, bacterial burden, and lung pathology were lower when mice were treated with amikacin, to which strain 0231 is sensitive (Fig. 6). Together these data validate the humanized dosing regimens of aztreonam and amikacin in this model of pulmonary P. aeruginosa infection mimics what is expected in the clinic, providing significant reduction in bacterial proliferation, dissemination, and lung pathology against *P. aeruginosa* that is sensitive to the antibiotic, but not bacteria that are resistant to the antibiotic.

**Fig. 5.**
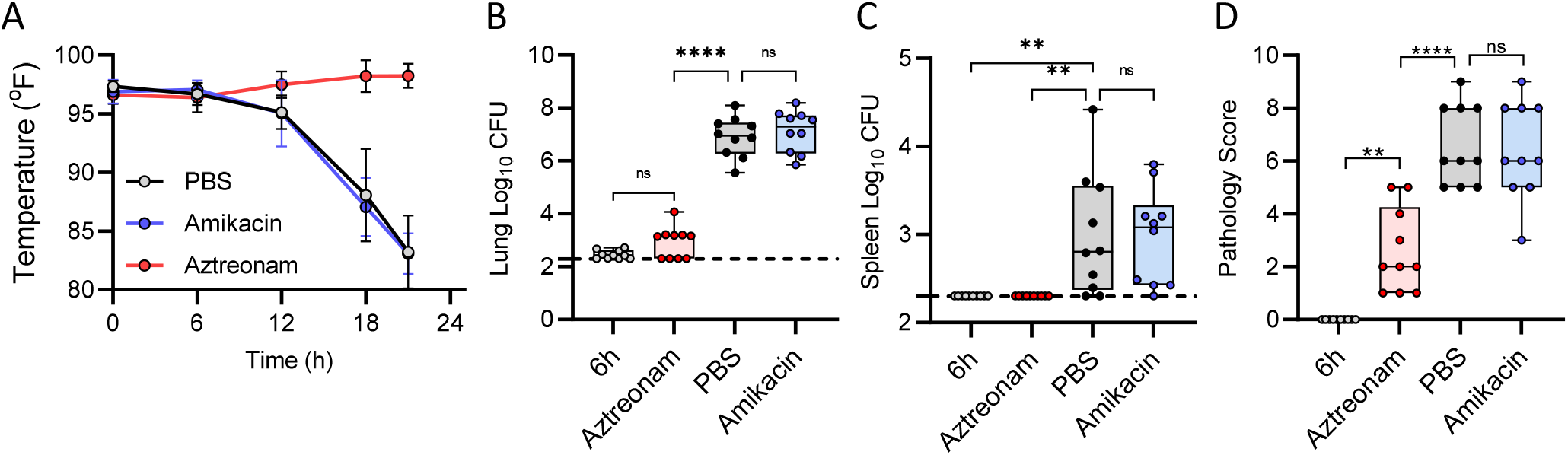
Affect of antibiotic treatment on *P. aeruginosa* 0230 infection. Equal numbers of male and female mice (n=10) were infected with 10x the LD_50_ of *P. aeruginosa* 0230. At 6 h animals were treated with PBS, aztreonam, or amikacin. (A) Temperature of animals during the course of the experiment. At 21 h mice were euthanized and bacterial numbers were enumerated from the (B) lungs or (C) spleens. (D) Lung pathology at 21 h. Data for B-D were compared to samples harvested from a group of animals at 6 h (prior to treatment) and each circle represents data from an individual mouse. The dotted line indicates the limit of detection. ANOVA w/ Tukey’s. **= p≤ 0.01; ****= p≤ 0.0001; ns= not significant.

**Fig. 6.**
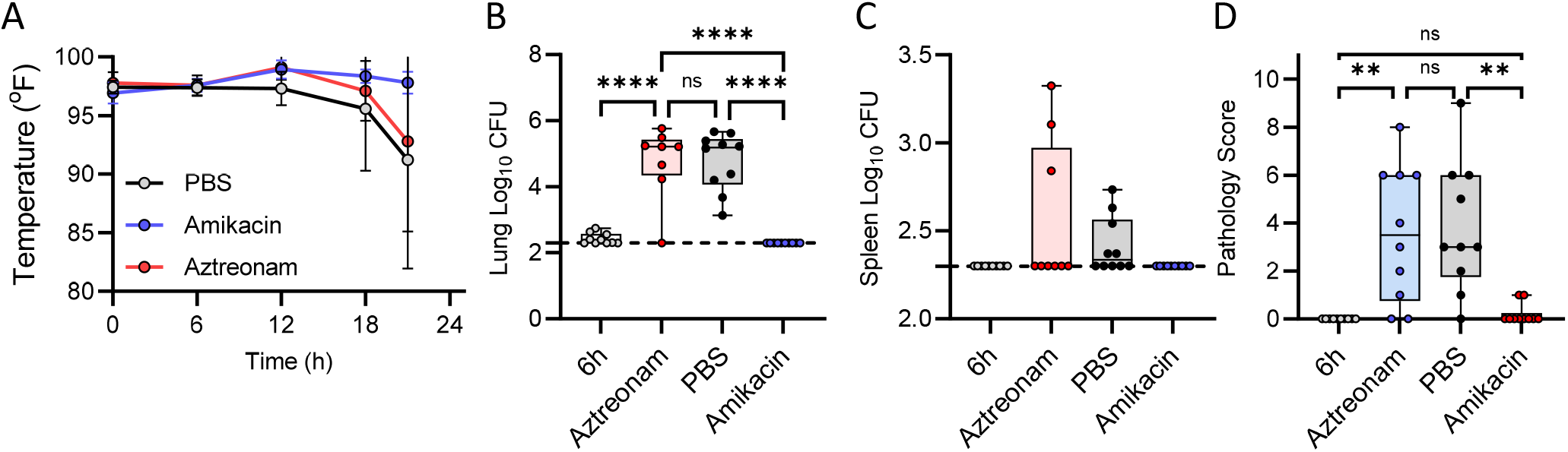
Affect of antibiotic treatment on *P. aeruginosa* 0231 infection. Equal numbers of male and female mice (n=10) were infected with 10x the LD_50_ of *P. aeruginosa* 0231. At 6 h animals were treated with PBS, aztreonam, or amikacin. (A) Temperature of animals during the course of the experiment. At 21 h mice were euthanized and bacterial numbers were enumerated from the (B) lungs or (C) spleens. (D) Lung pathology at 21 h. Data for B-D were compared to samples harvested from a group of animals at 6 h (prior to treatment) and each circle represents data from an individual mouse. The dotted line indicates the limit of detection. ANOVA w/ Tukey’s. **= p≤ 0.01; ****= p≤ 0.0001; ns= not significant.

### Validation of humanized doses of antibiotics – Long term survival

Having validated that humanized doses of aztreonam and amikacin can reduce bacterial burdens in infected mice based on antibiotic resistance profile, we next asked if these regimens could protect mice against lethal infection by *P. aeruginosa* that are sensitive to the antibiotic. Equal numbers of male and female mice were infected by direct instillation into the lungs with 10x the LD_50_ of *P. aeruginosa* strain 0230 (Azt^S^, Ami^R^) or 0231 (Azt^R^, Ami^S^). At 6 h post-infection, mice were administered 640 mg/kg aztreonam, 293 mg/kg amikacin, or vehicle only (PBS). Drug administration was repeated every 6 h for 5 days (120 h). At each administration, mouse temperatures were also measured. Mice were euthanized if they met predetermined endpoint criteria or at 5 days, and tissues were harvested for bacterial enumeration and histological analysis. As predicted by the significant reduction in bacterial burdens at 21 h post-infection (Fig. 5 and 6), mice receiving antibiotic did not display symptoms of infection and were protected from lethal infection, while mice receiving PBS all succumbed to infection within 48 h (Fig. 7 and 8). The only exception was one mouse in the amikacin group (Fig. 8). This mouse was also the only mouse that received antibiotics in which bacteria were recovered from tissues; bacterial burdens in all other antibiotic-treated animals were below our limit of detection. Minimal lung pathology was observed in antibiotic-treated animals, which was significantly lower than PBS-treated animals (Fig. 7E and 8E; p>0.001), with the exception of the one animal that succumbed to infection in the amikacin-treated group. As predicted by the bacterial burden within the lungs, this animal showed pathology similar to that observed in the PBS-treated cohort (Fig. 8E). Together, these data demonstrate that the antibiotic regimens determined by pharmacokinetic studies protected mice from lethal respiratory disease by *P. aeruginosa* that are sensitive to the antibiotic.

**Fig. 7.**
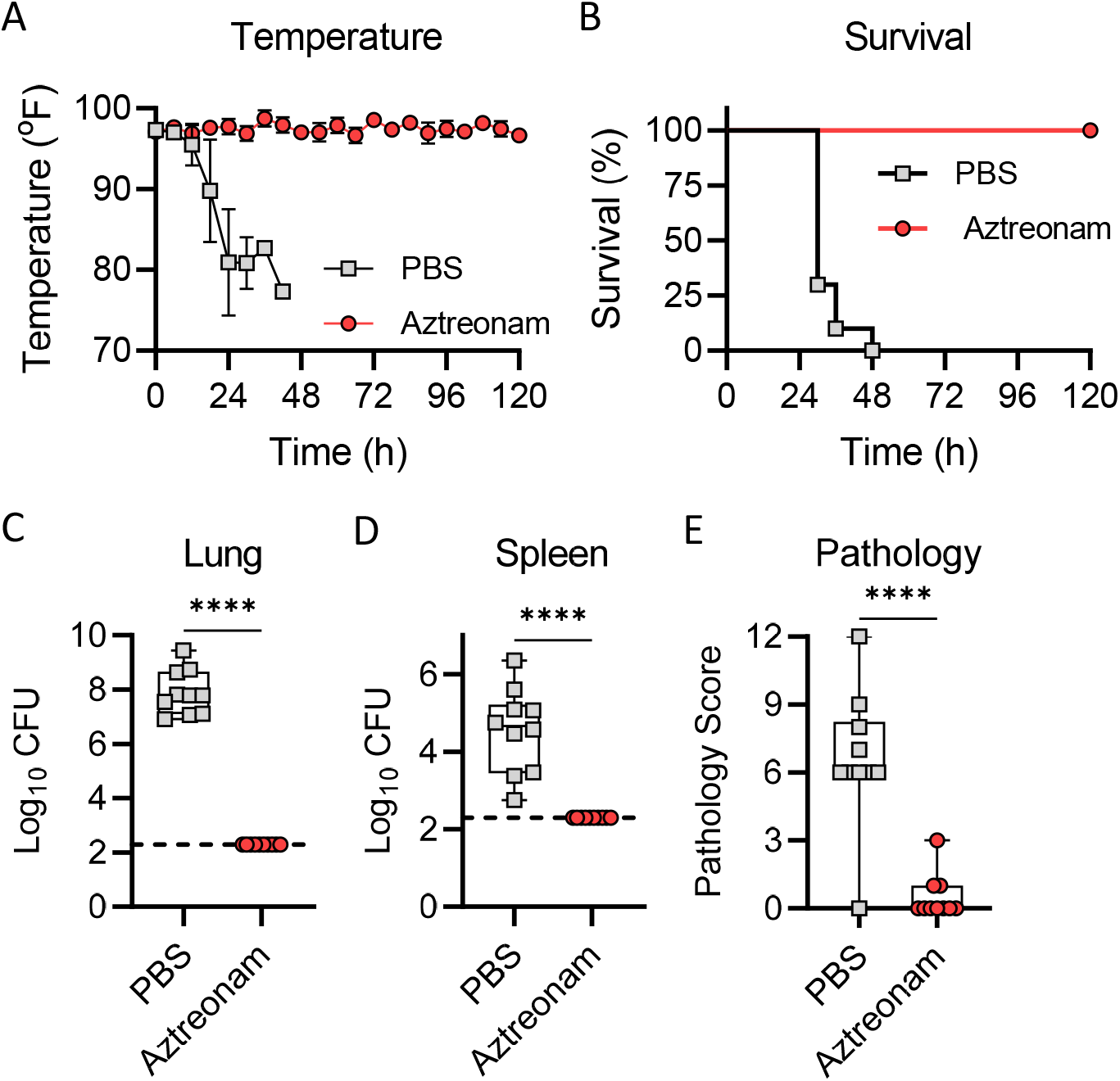
Aztreonam efficacy during 5-day treatment. Equal numbers of male and female mice (n=10) were infected with 10x the LD_50_ of indicated *P. aeruginosa* strain 0230. (A) Temperature measurements at indicated times. (B) Mouse survival. (C-E) Mice were euthanized if they reached predetermined endpoint criteria or at 5 days and bacteria were enumerated in the (C) lungs or (D) spleen. (E) A cross section of the lungs was excised, stained with H&E, and inflammation and pathology was assessed. For C-E, each symbol represents data from an individual mouse. Student’s t-test.****= p≤ 0.0001.

**Fig. 8.**
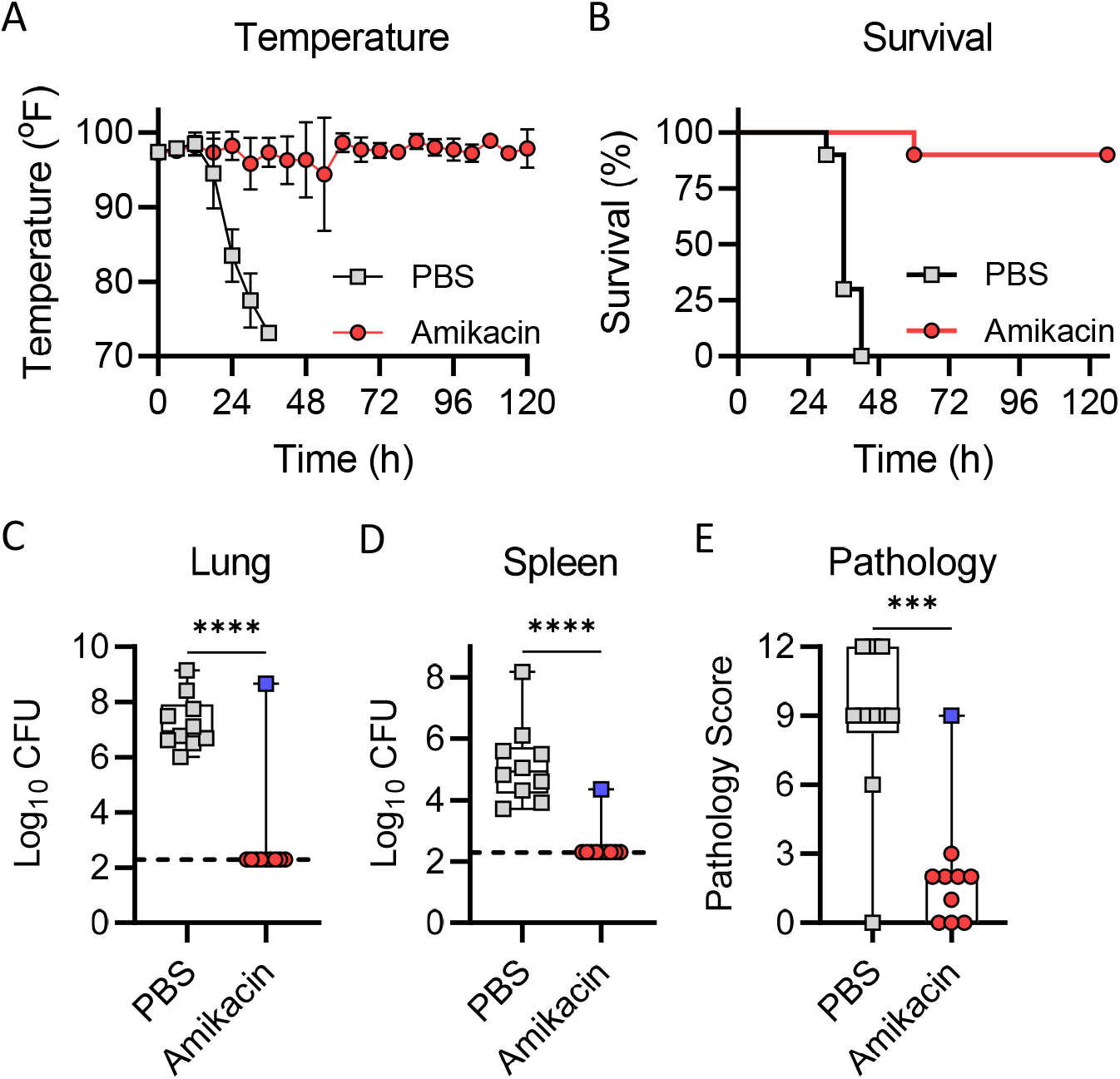
Amikacin efficacy during 5-day treatment. Equal numbers of male and female mice (n=10) were infected with 10x the LD_50_ of indicated *P. aeruginosa* strain 0231. (A) Temperature measurements at indicated times. (B) Mouse survival. (C-E) Mice were euthanized if they reached predetermined endpoint criteria or at 5 days and bacteria were enumerated in the (C) lungs or (D) spleen. (E) A cross section of the lungs was excised, stained with H&E, and inflammation and pathology was assessed. For C-E, each symbol represents data from an individual mouse. Blue square in the amikacin group indicates the mouse that succumbed to infection in this group. Student’s t-test. ***= p≤ 0.0001; ****= p≤ 0.0001.

## Discussion

The rise of antibiotic resistance in bacterial pathogens is far outpacing the discovery of new antimicrobials to treat these infections. Efforts to identify and test novel antimicrobials need to be implemented to ensure that we can treat pathogens that become resistant to all current therapies. In addition to research into the discovery of new antimicrobials, the development of robust preclinical models that can rapidly screen novel INDs and accurately predict clinical success are important to speed the drug development pipeline. Moreover, validated preclinical animal models can be used to support clinical studies in which patient population sizes may be small due to infrequency of infection by specific MDR bacteria. Therefore, our goal in these studies was to improve upon an existing murine model for the testing of antimicrobials against *P. aeruginosa* by expanding the number of strains of MDR *P. aeruginosa* that have been characterized for virulence in this model, developing humanized regiments of two conventional antibiotics that can be used as comparators in subsequent IND screening, and validating that the model can accurately differentiate between successful and unsuccessful treatments based on the inherent drug susceptibility of a given *P. aeruginosa* strain.

The model employed here is a transiently leukopenic murine model that best recapitulates acute pulmonary infection of immunocompromised patients. We previously demonstrated that the induction of leukopenia significantly reduces the dose of bacteria required to establish a lethal infection [9]. This reduction in infectious dose is important, as models requiring high doses of *P. aeruginosa* can artificially impact the interpretation of the efficacy of therapies, which we previously showed for meropenem using *P. aeruginosa* UNC-D. While treatment with meropenem could reduce bacterial burdens in an immunocompetent model, which required ∼10^8^ CFU to establish a lethal infection, the mice still succumbed to infection. In contrast, meropenem could reduce bacterial burdens and protect mice from lethal infection in the leukopenic model that only required ∼10^5^ CFU to establish a lethal infection [9]. However, the leukopenic model can still differentiate virulence between strains. We previously demonstrated this with two strains, the highly pathogenic lab adapted strain PA01 and a less virulent MDR clinical isolate UNC-D [9], and have now independently confirmed this using four diverse MDR strains. While the reasons for these differences in virulence have not been defined and are outside the scope of these studies, it highlights the variety of pathogenicity within clinical isolates of *P. aeruginosa*. Moreover, we hope that these data raise awareness for the need to fully characterize individual strains in animals to ensure proper infectious doses are used in subsequent in vivo testing of antimicrobials. Together, these two studies establish infection kinetics for six strains that are readily available through the NIH and CDC and increases the genetic diversity of the strains that can be used to test INDs in this preclinical model. This increased diversity, including in known antibiotic resistance mechanisms, more closely represents the clinical populations and should provide a better prediction of an IND’s success in the clinic.

The metabolism and physiology of mice differs from humans [18, 19]. These differences can have significant effects on many classes of drugs, resulting in drugs that demonstrate promise in the murine model but fail in clinical trials [20, 21]. However, because antimicrobials directly target the pathogen and not the host, positive results in the murine model have higher rates of successful translation in the clinic. Clinical success is further improved by incorporation of pharmacokinetic and pharmacodynamic analysis of antimicrobials in mice to identify driver indices that are independent of host metabolism [8]. Using aztreonam and amikacin, we established the feasibility of pharmacokinetics and pharmacodynamics in this model during infection and were able to use this data to establish a dosing regimen that closely mimics the parameters achieved during clinical treatment of humans. As such, we have shown that pharmacokinetics can be achieved with this model and applied during future characterization of INDs to establish the driver indices required for effective treatment during downstream clinical trials.

Finally, using aztreonam and amikacin we were able to validate that the model can clearly differentiate between successful and unsuccessful outcomes, which directly correlated with in vitro predictions of antibiotic susceptibility as expected. These data support that in vitro MICs for INDs should reliably justify whether a novel antimicrobial be further evaluated in the mouse model. Moreover, the characterization of these two antibiotics also provides robust comparators that can be included in future evaluations to help gauge relative success of novel INDs. Ideally, INDs would perform as well or better than these conventional antibiotics. Moreover, inclusion of aztreonam or amikacin during subsequent IND screening will provide data validating the model is performing reproducibly during each use, eliminating potential errant results by unforeseen circumstances.

In summary, the data presented here demonstrates a robust preclinical mouse model that can be used to 1) generate data to justify subsequent studies in larger animal models and IND applications; and 2) provide additional data to support the smaller datasets that are usually generated in clinical studies targeting rarer MDR bacterial infections.

## Materials and Methods

### Bacterial strains

The *P. aeruginosa* strains used in these studies were provided by the Center for Disease Control and Prevention as part of the CDC & FDA Antibiotic Resistance (AR) Isolate Bank. These strains have been sequenced and the antibiotic resistance profiles of each strain has been fully characterized (Table 1). Bacteria were routinely cultured on Luria-Bertani (LB) agar and LB-Lennox broth. For infection, bacteria were recovered from cryopreservation on LB agar prior to inoculation of LB-Lennox broth for overnight growth at 37°C with aeration. Bacteria were washed into 1x phosphate buffered saline (1xPBS) to the desired concentration based on OD_600_-based estimates. All bacterial inoculums were confirmed by serial dilution and enumeration on LB agar.

### Animals

All animal studies were approved by the University of Louisville Institutional Animal Care and Use Committee (IACUC no. 18368). Six-week-old male and female BALB/cJ mice (Jackson Laboratory, Bar Harbor, ME) were acclimated on site for eight days prior to bacterial challenge. Upon arrival, a temperature transponder (Biomedic Data Systems; Seaford, DE) was implanted subcutaneously at the scruff of the neck. Neutropenia was induced in the animals via intraperitoneal injection of cyclophosphamide as previously described [9]. Neutropenia (>90% depletion) was confirmed by complete blood cell counts one day prior to infection using a Hemavet 950 (Drew Scientific, Miami Lakes, FL). Group sizes for all studies were 10 animals, consisting of 5 males and 5 females.

### Respiratory Challenge with *P. aeruginosa*

Mice were challenged with saline suspensions of *P. aeruginosa* by intubation-mediated intratracheal inoculation (IMIT) as previously described [9, 14]. Animals were monitored for the development of moribund disease every 8h after infection. Animals that met predetermined endpoint criteria (body temperature ≤ 80.5°F or loss of righting reflex) were humanely euthanized by carbon dioxide asphyxiation and scored as succumbing to infection 8h later.

### Natural history of *P. aeruginosa* infection

Upon euthanasia, lungs and spleen were harvested. A representative section of lung was excised, fixed in 10% neutral buffered formalin for 24 h, and transferred into 70% ethanol prior to hematoxylin and eosin (H&E) staining. Stained sections were scored by a Board-Certified Veterinary Pathologist at the Iowa State University Veterinary Pathology Comparative Pathology Core and compared to naive control lungs. Pathology scoring was made on a four-point, four criteria system with a maximum score of 16 points. Tissues were scored in the areas of inflammation, infiltrate, necrosis, and other (including hemorrhage), and points assigned as: no significant finding (0), minimal (1), mild (2), moderate (3) and severe (4) pathology. Bacterial were enumerated from remaining lung tissue and spleens as previous described [9].

### Defining human simulated doses of antibiotics

Single-dose plasma pharmacokinetic studies were performed in infected mice. For each drug, four doses were utilized to generate a robust pharmacokinetic dataset from which a humanized dosing regimen was determined. Groups of four animals were administered single subcutaneous doses of antibiotic (Amikacin 320, 640, 960, and 1280 mg/kg; Aztreonam 320, 640, 1280, 2560 mg/kg). At six defined time points (5, 10, 15, 30, 60, and 120 min post-administration of the antibiotic), animals were euthanized and plasma collected from each mouse. Plasma samples were stored at -80°C until antibiotic quantification. Antibiotic concentrations in plasma were quantified and analytically assessed by high performance liquid chromatography with an Agilent Compact 1260 HPLC system as previously described [22, 23]. Briefly, aztreonam was analyzed with a XBridge BEH C18 column and VanGuard Cartridge. An isocratic flow rate of 0.5 ml/min of the eluent (0.005 M tetrabutylammonium hydrogen sulfate [pH 3.0]- acetonitrile [volume ratio=88:12]) and UV 280 nm were used for the analysis. Validation range and performance were recorded as: LLOQ:5 μg/ml; ULOQ: 250 μg/ml; QC: 25 μg/ml; Precision:96.36%; Accuracy: 88.26%. Amikacin was conjugated with 4-chloro-3,5-dinitrobenzotrifluoride (CNBF) before analysis. The resulting Amikacin-CNBF was analyzed with an Agilent Eclipse Plus C18 column. A gradient of methanol and DI water (0.1% TFA) were used, starting with methanol percentage increasing from 35% to 60% in the initial 2 min, then 60% of methanol was kept for 1.5 min before further increasing to 90% in the next 1 min, followed with another 1 min of 90% methanol. The percentage of methanol was then reduced to 35% in 2 min and kept at 35% for another 1.5 min. A flow rate of 1.6 ml/min and UV 238 nm were used for the analysis. Validation range and performance were recorded as: LLOQ: 71 μg/ml; ULOQ: 1428 μg/ml; QC: 357 μg/ml; Precision: 94.98%; Accuracy: 95.59 %.

Pharmacokinetic parameters (means ± standard deviations) were calculated using a time ordered dataset generated from the pooled PK data. Elimination half-life (t1/2), AUC_0–∞_, and Cmax, were calculated using a noncompartmental model with mean concentration values from each group of mice. The half-life was determined by linear least-squares regression. The AUC was calculated from the mean concentrations using the trapezoidal rule. Pharmacokinetic estimates for dose levels that were not directly measured were calculated using linear interpolation for dose levels between those with measured kinetics and linear extrapolation for dose levels above or below the highest and lowest dose levels with kinetic measurements.

Human pharmacokinetic data for aztreonam and amikacin was utilized to generate simulated human drug concentration-time exposure profiles for intravenous aztreonam 2 g every 8 h and amikacin 15 mg/kg once daily [15-17]. Using the pharmacokinetic results generated above, we modeled dosing regimens in the mice that would be expected to provide similar drug concentration-time exposures to that in humans. We utilized the PK/PD parameter of time drug concentration exceeds the MIC over the dosing period (%T>MIC) for aztreonam and evaluated both the maximal concentration relative to MIC (C_max_/MIC) and the 24 hour area under the drug concentration curve relative to MIC (AUC_0-24_/MIC) for amikacin [24].

### Evaluation of antibiotic efficacy: Reduction in bacterial burdens

Six h after IMIT instillation of *P. aeruginosa* 0230 or 0231 (10x the LD_50_), mice were administered humanized doses of either amikacin (293 mg/kg Q6h) or aztreonam (640 mg/kg Q6h). Animal temperatures were also acquired at every treatment. 21 h post-infection, mice were euthanized, and bacterial numbers were determined in the lungs and spleen by serial dilution and enumeration on agar plates. A section of the lungs was also processed for histological analysis as described above.

### Evaluation of antibiotic efficacy: Overall survival

Six h after IMIT instillation of *P. aeruginosa* 0230 or 0231 (10x the LD_50_), mice were administered humanized doses of either amikacin (293 mg/kg Q6h) or aztreonam (640 mg/kg Q6h) for 5 days. Animal temperatures were also acquired at every treatment. Blood, lungs, and spleens were harvested from moribund animals or at 5 days post-infection from animals that survived to the end of the study. Bacterial numbers in these tissues were determined by serial dilution and enumeration on agar plates. A section of the lungs was also processed for histological analysis as described above.

### Statistical analysis

Kaplan-Meier survival curves were fit to the data for each bacterial strain and the median survival time noted by gender and overall. The LD_50_ was calculated by fitting the logistic model (%dead = log_10_[dose]) and then calculating the dose in which 50% of the mice are predicted to survive. To assess the natural history of infection, a repeated measures linear model was fit to temperatures by bacterial strain assuming an autoregressive correlation structure. Power calculations estimated the significant lung burden (log_10_) to be seen for 80% power and sample sizes of 10 mice per treatment groups assuming an alpha of 0.05 and a possible 20% increase in variability within the treatment group. The disease burden on the lungs and spleen were modeled on the log_10_ scale with sex and hour as the independent variables. Fisher’s Exact tests compared the incidence of pathology. All data were analyzed by sex and in aggregate. No significant differences by sex were observed.

## Acknowledgments

This work was supported by funding from the FDA, contract HHSF223201810171C (MBL). We thank members of the FDA advisory panel for their advice during these studies.

